# Coupled Reaction Networks for Noise Suppression

**DOI:** 10.1101/440453

**Authors:** Fangzhou Xiao, Meichen Fang, Jiawei Yan, John C. Doyle

## Abstract

Noise is intrinsic to many important regulatory processes in living cells, and often forms obstacles to be overcome for reliable biological functions. However, due to stochastic birth and death events of all components in biomolecular systems, suppression of noise of one component by another is fundamentally hard and costly. Quantitatively, a widely-cited severe lower bound on noise suppression in biomolecular systems was established by Lestas *et. al.* in 2010, assuming that the plant and the controller have separate birth and death reactions. This makes the precision observed in several biological phenomena, e.g., cell fate decision making and cell cycle time ordering, seem impossible. We demonstrate that coupling, a mechanism widely observed in biology, could suppress noise lower than the bound of Lestas *et. al.* with moderate energy cost. Furthermore, we systematically investigate the coupling mechanism in all two-node reaction networks, showing that negative feedback suppresses noise better than incoherent feedforward achitectures, coupled systems have less noise than their decoupled version for a large class of networks, and coupling has its own fundamental limitations in noise suppression. Results in this work have implications for noise suppression in biological control and provide insight for a new efficient mechanism of noise suppression in biology.

## I. Introduction

Many important processes in living cells, such as gene expression, are intrinsically stochastic [1]. The effect of noise is further enhanced by several biological factors, such as the low copy number of many important molecules including DNA, and the fact that even controllers performing noise suppression consists of intrinsically stochastic molecular components themselves [1], [2], [3]. In particular, assuming that the controller and the plant dynamics are separate biochemical processes, Lestas *et. al.* [2] showed that there is a severe lower bound on the noise for the plant component of a chemical reaction network due to the intrinsic stochasticity of controller components in chemical reactions. Quantitatively, it states that the lower bound for noise is typically inversely proportional to the quartic root of the signaling rate, in contrast with a square root result that is familiar in electrical engineering and statistics. This bound implies that significant noise is inevitable, despite the regulatory mechanisms found in biology that could suppress noise [4].

Specifically, the bound implies that noise suppression in cells is usually prohibitively expensive, as reducing noise by 10 fold would require 10,000 fold increase in signaling rate, which corresponds to 10,000 fold faster production and degradation of molecules.

On the other hand, we observe almost deterministic precision in many biological processes, such as the synchronization of molecular oscillations in mammalian circadian clock [5], precise timing of gene expression dynamics in cell cycle [6] and the cell type differentiation through gene expression patterns during development [7]. This observation naturally urges one to seek biologically relevant low-cost noise suppression mechanisms that are beyond the scope of systems investigated in [2].

In this work we propose coupling as such a noise suppression mechanism that is omnipresent in biological processes. Coupling is used here to describe chemical reactions where the number of more than one chemical species are altered simultaneously, e.g., reactions converting one species into another one, or those producing one molecule each of two species simultaneously. Hence, coupling naturally goes beyond the assumption in [2] that the plant and the controller dynamics are separate biochemical processes, since the increase and/or decrease of a plant component and a controller component could be coupled in a reaction once we allow coupling.

Many natural and genetically engineered circuits in biology have a coupling interpretation. In bacteria, genes are commonly grouped into operons, so they are transcribed and regulated together [8]. Non-coding RNAs with regulatory functions such as microRNA and siRNA are commonly or designed to be in the same transcript with the mRNA they regulate [9], [10]. In metabolic networks, signaling cascades and allosteric regulation, one protein may switch from an inactivated state to an activated state by a reaction which may be regulated by the protein or its downstream products [8], [11]. Regulatory functions in synthetic biology are commonly implemented in binding proteins linked with functional domains, where the link is cut to transform the protein-domain complex into a protein and a domain separately, which could have different regulatory properties than the complex [12], [13].

The fact that specific coupling mechanisms, e.g. putting genes on the same operon, could influence the system’s performance is commonly known. Coupling as a noise suppression mechanism is also rather intuitive from an information-theoretic point of view. Indeed, earlier studies [10], [14], [15], [16] have shown a few specific examples of coupled reactions could have less noise than the decoupled versions.

However, it is not known what are the general conditions for coupling to suppress noise, the fundamental limitations on noise suppression once we allow coupling (in the spirit of [2]), or how to use coupling when designing biochemical network architectures.

In this work we set off to provide some initial answers to these problems. After reviewing the notions of chemical reaction networks and methods to analyze them in Section II, we show how coupling could suppress noise through a simple example in Section III-A. We then demonstrate the power of coupling via a biologically plausible coupled system that suppresses noise below the lower bound of Lestas *et. al.* [2] in Section III-B. Following Lestas *et. al.* [2], we establish a similar bound for coupled reaction networks in Section III-C to better understand how it compares with decoupled systems and when is coupling most useful. Lastly, in Section III-D, we investigate all possible two-node chemical reaction network architectures to gain insight on the advantage of coupled networks versus decoupled ones, feedback versus feedforward architectures, limitations of coupling, and how to choose network architectures to maximize coupling’s effect on noise suppression.

## II. Background

### A. Chemical Reaction Networks

Here we provide a minimum background on stochastic chemical reaction networks (CRNs) and relevant analysis methods. For more details, see [3], [17], [18].

A chemical reaction network (CRN) is a collection of reactions of the following form:

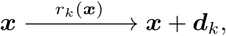

where 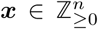 is the state vector describing the number of molecules of the *n* chemical species in the system, *k* = 1, *…, m* indexes *m* reactions, and *r*_*k*_ : ℝ^*n*^ → ℝ_≥0_ is the propensity function which describes the rate of reaction *k*. Technically, the system dynamics is described as a continuous-time Markov jump process with 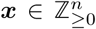 as states of the system, and *r*_k_ (***x***)Δ*t* describes the probability that, if the system is in state ***x*** at time *t*, it will jump to state ***x*** + ***d***_*k*_ within time Δ*t*, for small Δ*t*. Note that even though the mass action law, where *r*_*k*_ are always a special form of polynomial of ***x***, is commonly assumed [19], we do not make that assumption here.

The stochastic description of the system dynamics is described in terms of the chemical master equation (CME), which describes the evolution of the probability distribution for molecular counts ***x*** [20]:

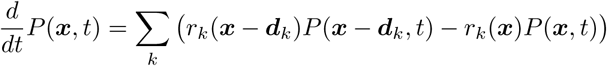

where *P* (***x***, *t*) is the probability that the system has ***x*** number of molecules at time *t*.

For the mean, we have

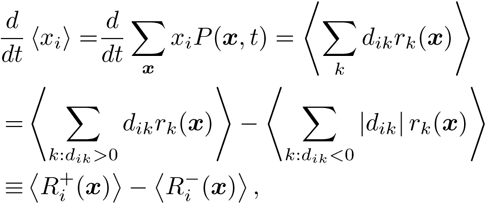

where the last equality was used as the definition for 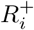 and 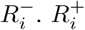, the birth flux of *x*_*i*_, is the summed rate of all reactions that increases *x*_*i*_, while 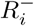, the death flux, is the summed rate of all reactions that decreases *x*_*i*_. Note that, at steady state, we have 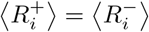.

It is worth noting that, in the limit where the reaction volume is large while the number of molecules per volume remains finite (made precise in [21]), the system dynamics can be described by a deterministic rate equation:

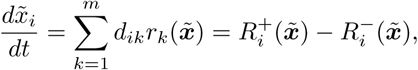

where 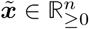 is the continuous concentrations of chemical species, instead of discrete molecular counts 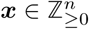.

Through similar calculations as in the equation for the mean, we see that, at steady state, we have the following equation for covariance:

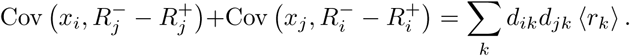

One major achievement of the theoretical investigations in [3] is the re-writing of the above equation using physically interpretable quantities. There-written equation is as follows:

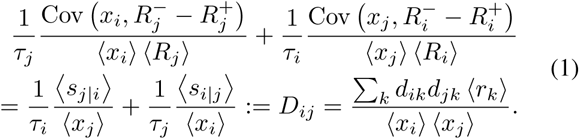

Here *τ*_*i*_, the average life time of an *x*_*i*_ molecule, and ⟨*s*_*i*|*j*_ ⟩, the average step sizes or co-step sizes, are introduced. As these are key concepts utilized for this work, they are explained in detail below.

#### Step sizes

For *i* = *j*, we define the average step size of *x*_*i*_ as ⟨*s*_*i*|*i*_⟩*=* Σ_*k*_ |*d*_*ik*_| *ρ*_*ik*_, where 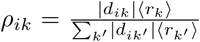, the probability that when *x*_*i*_ changes, that change comes from reaction *k*. The notation ⟨*s*_*i*|*i*_⟩ signifies that this is average step size of *x*_*i*_ conditioning on that *x*_*i*_ changes. For example, if *x*_1_ has only one birth reaction *x*_1_ → *x*_1_ + 1 and one death reaction *x*_1_ → *x*_1_ *-* 10, then 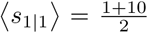, because birth and death fluxes are always equal at steady state.

For *i* ≠ *j*, the co-step size is defined as ⟨*s*_*j*|*i*_⟩ *=* Σ_*k*_ *ρ*_*ik*_ |*d*_*jk*_| sgn {*d*_*ik*_*d*_*jk*_}. So ⟨*s*_*j*|*i*_⟩ is the average change of species *x*_*j*_ conditioning on that *x*_*i*_ changes simultaneously in these reactions. Note that this could be positive or negative. From the definition, we see that the co-step size ⟨*s*_*i*|*j*_ ⟩captures whether birth and death of *x*_*i*_ and *x*_*j*_ are coupled in some reaction. For example, if the only reaction that have simultaneous changes to *x*_1_ and *x*_2_ is (*x*_1_, *x*_2_) → (*x*_1_ *-* 1, *x*_2_ + 1), i.e. one *x*_1_ becomes one *x*_2_, and *x*_2_ have no other birth reactions, then 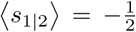. It is negative because when *x*_1_ decreases, *x*_2_ increases. It is divided by 2 because this reaction accounts for all of *x*_2_’s birth, therefore half of *x*_2_’s changes.

Note that the co-step sizes and the birth and death fluxes are related to each other: ⟨*s*_*j*|*i*_ ⟩ ⟨*R*_*i*_⟩ = ⟨*s*_*i*|*j*_⟩ ⟨*R*_*j*_⟩.

#### Life times

For general stationary stochastic processes describing the increase and decrease of a quantity, e.g. molecule count, we can define the average life time as the average time that one molecule lasts before its degradation. Little’s law in queueing theory then states that the average life time is equal to the ratio between the average number of such molecules and its average birth or death rate, regardless of how nonlinear the reaction rates are. This could be intuitively understood through the following non-rigorous calculation, where 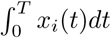 is the sum of life time of all *x*_*i*_ molecules in *T*, and 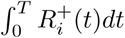 is the total number of *x*_*i*_ molecules born in *T* :

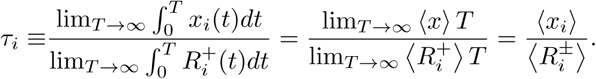

Therefore, we have 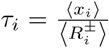, the average life time or the time scale of *x*_*i*_.

### B. Linear Noise Approximation

We see that the covariance equation (1) involves terms of the form Cov 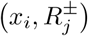, which would involve higher order moments if 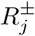 is nonlinear. The resulting system of moment equations is generally infinite dimensional and hard to solve. This naturally calls for moment closure methods to approximately solve this system [22]. One particularly simple and analytically tractable method is linear noise approximation (LNA). LNA could be derived by assuming that the mean ⟨***x*** ⟩ is close to a fixed point so that 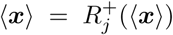, and by simply approximating birth and death rates by their first order Taylor expansion: 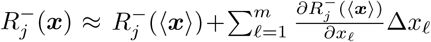. Define the logarithmic gains 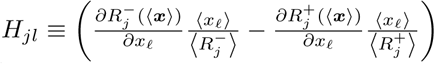 [17], equation (1) becomes

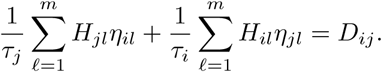

In matrix form, we have

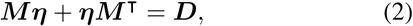

where 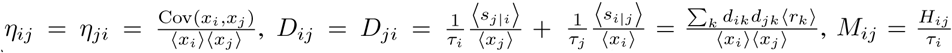. Note that an internal relation needs to be utilized when solving this system of equations:

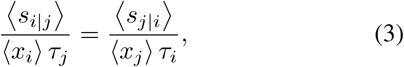

which is derived from the relation ⟨*s*_*i*|*j*_ ⟩ ⟨*R*_*j*_⟩ = ⟨*s*_*i*|*j*_ ⟩ ⟨*R*_*i*_⟩.

Equation (2) is called the fluctuation dissipation theorem in the statistical physics community [23] and Lyapunov equation in the control community [24]. Note that the normalized covariance ***η*** is a solution to this Lyapunov equation shows that it is finite if and only if the linearized deterministic system 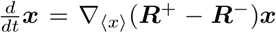 is asymptotically stable, i.e. *-* ***M*** is Hurwitz. Hence there exists a general relationship between deterministic stability and stochastic variance [25].

As LNA is the main tool of analysis in this work, more comments on the effectiveness of LNA is in order. Although LNA was historically derived as a second order system size approximation of the chemical master equation, resulting in a Fokker-Planck equation with a stochastic differential equation interpretation and a Gaussian steady state distribution [26], this is not necessary. LNA, as well as higher order approximations, could be derived as Taylor expansions on the moment equations [27]. In fact, it could be shown that LNA could be interpreted as a linear propensity CRN approximation that preserves discreteness and non-negativity of the state variables (see Section V-B). So violating discreteness and non-negativity are not valid grounds for rejecting the usefulness of LNA. LNA fails just like how deterministic linearization fails: when nonlinearity significantly influences the state trajectory.

## III. Results

### A. Coupling could reduce noise

Here we illustrate how coupling could reduce noise through one simple example with very explicit analysis.

Consider the following feedforward network, called a “sniffer” system in biological literature [28].

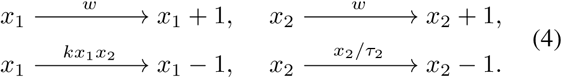

It consists of two components *x*_1_ and *x*_2_, both activated by an external input *w*, while *x*_2_ acts as an enzyme that catalyze the degradation of *x*_1_. The system (4) does not couple the birth events of *x*_1_ and *x*_2_, so they are produced with rate *w* through separate reactions. The corresponding coupled version of system (4) keeps the last two death reactions the same while substituting the following reaction for the first two birth reactions:

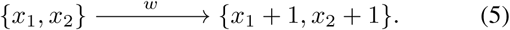

For both coupled and decoupled versions, we see that the birth rates of *x*_1_ and *x*_2_ are 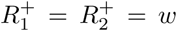, their death rates are 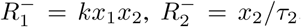, and their step sizes are ⟨*s*_11_ ⟩ = ⟨*s*_22_ ⟩ = 1. For the coupled version, the co-step sizes are 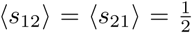, indicating half of the flux are through a coupled reaction where *x*_1_ and *x*_2_ are increased or decreased with the same number, while both co-stepsizes are 0 for the decoupled version. Applying LNA (see Section II), the steady states are ⟨*x*_1_ ⟩ = *τ*_2_/*k*, ⟨*x*_2_ ⟩ = *w*/*τ*_2_ and the time scales are 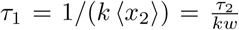. The gain matrix is 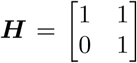. The diffusion matrix ***D*** in equation (2) are the following:

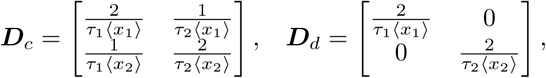

where ***D***_*c*_ is for the coupled case and ***D***_*d*_ is for the decoupled case.

Solving equation (2) with ***D*** = ***D***_*d*_ for the decoupled case, we have

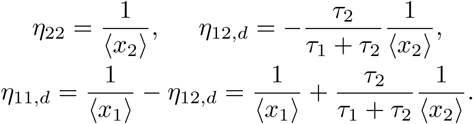

We see that the noise of *x*_1_ in the decoupled case, *η*_11,*d*_, can be decomposed into two parts: the first t erm 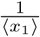 that is the intrinsic noise of *x*_1_ doing birth-death by itself, and the second term that is the noise of *x*_2_ carried over to *x*_1_ through its regulation on *x*_1_.

In comparison, solving equation (2) with ***D*** = ***D***_*c*_ for the coupled case, with the additional constraint *τ*_2_ ⟨*x*_1_⟩= *τ*_1_ ⟨*x*_2_⟩ which is a consequence of equation (3), we have the following:

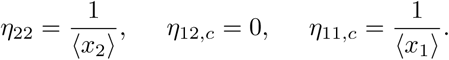

Note that *η*_11,*c*_ is the same as that of a simple birth-death process of *x*_1_ by itself, with no interaction with *x*_2_ at all. The noise for the decoupled case is larger than the coupled case by 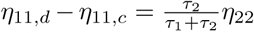, *x*_2_’s noise passed onto *x*_1_ through their interaction. This indicates that the noise contributed to *x*_1_ from the interaction with *x*_2_ in the decoupled case is cancelled out here due to coupling.

This example shows that coupling could indeed reduce noise. In fact, similar circuits have been analyzed in [14], [10] that report specific RNA circuit designs with coupling could suppress gene expression noise.

### B. Coupling suppresses noise below Lestas et. al. bound

We show here through a biologically plausible example that coupled systems could have noise below the bound described in [2].

Consider the following system:

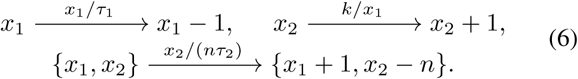

Here *n* molecules of *x*_2_ can be converted to 1 molecule of *x*_1_, resulting in a negative coupled reaction. Besides this, *x*_1_ has first order death reaction by itself, while *x* _2_ is produced through a reaction that is suppressed by *x*_1_.

When *n* = 1, the coupled reaction could be an enzyme switching from an inactive state *x*_2_ into an active state *x*_1_, which in turn suppresses the production of this enzyme *x*_2_. Alternatively, it could be a binding protein in complex with a functional domain *x*_2_ being digested into separate parts, where the free binding protein *x*_1_ suppresses the production of this complex *x*_2_. When *n >* 1, the coupled reaction could be protein subunits *x*_2_ forming a protein complex *x*_1_ involving *n* such subunits (e.g. *n* = 2 corresponds to a dimer), while *x*_1_ has one of its active functions the suppression of subunit *x*_2_’s production. Therefore, this system is biologically plausible with potential straight-forward implementations through tools in synthetic biology.

To compare with the Lestas *et. al.* bound, we need to delve into the details of their work [2]. Lestas *et. al.* considered reaction networks of the following form:

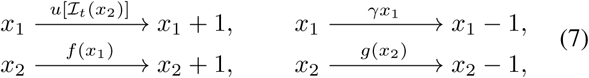

where *x*_1_ is the plant species whose noise is to be controlled, while *x*_2_ is the signaling controller species whose information on *x*_1_ is the only source of information on *x*_1_ we could use to suppress its noise. The signaling of *x*_2_ is through its birth events, as the birth rate *f* is dependent on *x*_1_. For example, if *f* (*x*_1_) = *x*_1_, so *x*_1_ catalyzes the production of *x*_2_, then we would estimate *x*_1_ to be large if we observe a high density of *x*_2_’s birth events. Therefore, information about *x*_1_’s abundance could be extracted through a trajectory of *x*_2_’s birth events. Because *x*_2_’s death rate *g* does not depend on *x*_1_, the death events does not give any information about *x*_1_. So we ignore death events and focus on *x*_2_’s birth events. On the other hand, with the information about *x*_1_ extracted through observations on *x*_2_, we try to suppress *x*_1_’s noise by controlling its (possibly non-Markovian) production rate, *u*[ℐ_*t*_(*x*_2_)], which could be an arbitrary non-anticipatory functional allowing dependence of all the information ℐ_*t*_(*x*_2_) of past *x*_2_. In control theory terms, *x*_1_ is the plant, birth events of *x*_2_ with signaling rate *f* is the sensor, and *x*_1_’s birth rate *u* is the controller actuation.

Lestas *et. al.* then proceeded to use sensor and actuator separation in order to first bound the channel capacity in terms of *f* and then bound the noise in terms of channel capacity. The bounds are the following:

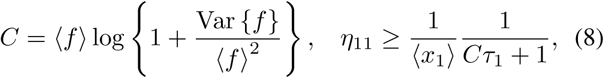

where *C* is an achievable upper bound for the channel capacity of *x*_2_’s birth events assuming finite mean and variance of *f*, and *τ*_1_ is the average life time of *x*_1_ as defined in Section II, which is equal to 1*/γ* for system (7). If we can write the mean and variance of *f* in terms of those of *x*_1_ (e.g. when *f* ∝ *x*_1_), then these two bounds could be combined to produce a lower bound of *x*_1_ in terms of only *x*_1_’s mean, variance, and time scales. Importantly, when *f* is a linear function of *x*_1_, i.e., *f* = *αx*_1_, this bound is:

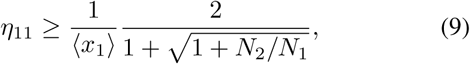

where *N*_1_ = ⟨*x*_1_⟩ and *N*_2_ = *α* ⟨*x*_1_⟩*τ*_1_ are the effective signaling rates, defined as the number of birth events of *x*_1_ and *x*_2_ during an average life time of *x*_1_, i.e., *τ*_1_, respectively. This is the quartic bound for the coefficient of variation 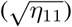 mentioned in Section I.

Going back to our example (6), we see that the number of *x*_2_ molecules consumed in the coupled reaction, *n*, is equal to the relative signaling rate 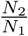 as defined above. So we expect to see better noise suppression with increasing *n*. The birth rate satisfies *f* = *k/x*_1_, which is not a linear function of *x*_1_, so we need to use equation (8) instead of the linear bound (9). In simulation, we need to estimate mean and variance of *f* to calculate *C*, which in turn enables calculation for lower bound of *η*_11_ in equation (8). It is important to note that we violated none of the assumptions about (7) in Lestas *et. al.* bound (8) except allowing coupling among reactions.

Using LNA, we could analytically solve system (6) to have 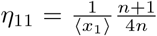 in the fast controller dynamics limit, where *τ*_2_ ≪ *τ*_*1*_ (see Section III-D). The simulation result comparing the simulated noise of *x*_1_, its LNA analysis, and the Lestas *et. al.* bound in equation (8) is shown in Figure 1. We see that the noise is indeed well below the theoretical bound of Lestas *et. al.* for not too large *n*, which is the more biologically plausible parameter regime.

**Fig. 1.**
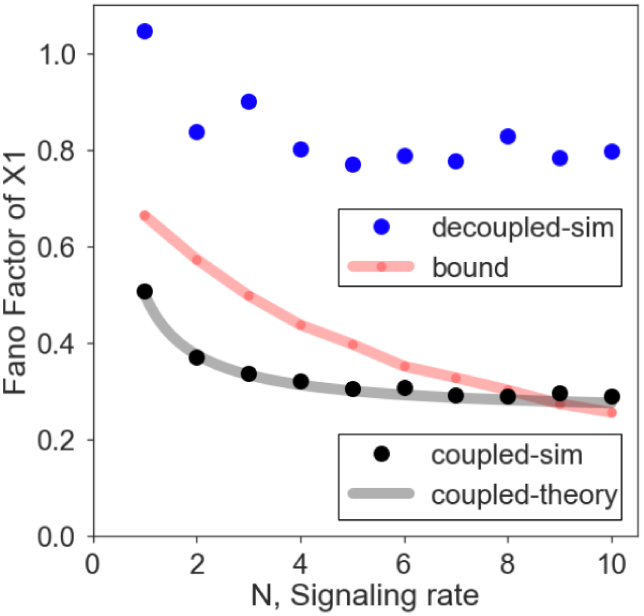
Noise of example (6) in Section III-B compared to Lestas *et. al.* bound in equation (8) [2]. The *y* axis is the variance divided by mean of *x*_1_, or its Fano factor. The *x* axis is the signaling rate *N* = *N*_2_*/N*_1_, taking integer values from 1 to 10. The Gillespie simulation results of system (6) are the black dots (coupled-sim), in good agreement with the theoretical results based on LNA shown as the light grey curve (coupled-theory). The theoretical bound of equation (8) with estimated capacity is shown as the light red curve (bound). The same simulation results for the decoupled version of system (10) are the blue dots (decoupled-sim). Parameters used for simulation: *k* = 10000,*τ*_1_ = 100,*τ*_2_ = 1.

We also did simulation for the decoupled version of system (6), where the following two reactions substituted the coupled reaction:

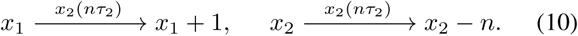

The simulation results for this decoupled version are the blue dots in Figure 1, showing that it is indeed above the Lestas *et. al.* bound, as the decoupled case satisfied all of the assumptions in their work [2].

It should also be noted that although this specific example (6) could achieve noise below the bound of [2], the scaling of noise in *N* in this system is actually worse than quartic root because it has a lower bound of 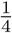, as is shown by the black curve’s convergence to 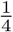 in Figure 1 as well as the LNA calculation. We will see in Section III-D that there exists in general a non-zero lower bound on the part of noise that can be suppressed through coupling. In other words, once the architecture of a system is determined, there is a lower bound on how much noise can be reduced by coupling. For a direct comparison, in the following section, we establish a lower bound on noise suppression when coupling reactions are allowed using information theoretic methods in [2].

### C. Bound on noise suppression with coupling

To better understand how coupling changes the fundamental limit on noise suppression, here we establish a bound similar to (9) that allows for coupled reactions. For detailed derivation, see V-C.

Let us first understand how coupled reactions go beyond bound (9). Bound (9) assumes that the birth events of *x*_2_ is a Poisson channel whose rate is *x*_2_’s birth rates. However, when the birth or death events of *x*_1_ and *x*_2_ are coupled, the mutual information between the two increases, since a change of *x*_2_ due to this coupled reaction always comes with a known change of *x*_1_ at exactly the same time. Hence the key to establish a bound that includes coupling is in bounding the mutual information through this coupled birth-death.

Consider a coupled reaction in place of the decoupled *x*_2_ *→ x*_2_ + 1 in equation (7):

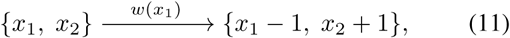

which could be regarded a simple conversion reaction where one *x*_1_ is converted into one *x*_2_. We could split this channel into two Poisson channels, where one is *x*_2_*→x*_2_ + 1 with rate *w*(*x*_1_) just like the decoupled case, and one is *x*_1_ *→ x*_1_ *-*1 with rate that is a function of *x*_1_ and *x*_2_, such that it becomes infinitely fast whenever it sees *x*_2_ change so as to bring *x*_1_ to match the change of *x*_2_, and once *x*_1_ changed it becomes 0. In other words, the *x*_1_*→x*_1_ *-*1 reaction is to track the change of *x*_2_ instantaneously.

With this consideration, we could bound the channel capacity of reaction (11) by ⟨*w*⟩ ln(1+1*/* ⟨*w*⟩)+Var {*w*}*/* ⟨*w*⟩. The first term comes from using equation (8) to bound the channel capacity of the *x*_2_ → *x*_2_+1 reaction with rate *w*(*x*_1_). The second term comes from the infinitely fast reaction *x*_1_*→x*_1_*-*1 mimicing the coupling. If we assume first order reactions *w*(*x*_1_) = *βx*_1_, then we can derive a bound in a form similar to Lestas *et. al.* bound in equation (9):

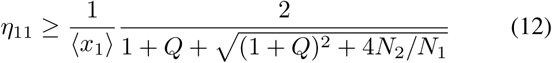

where *Q* = *N*_2_ ln(1 + *τ*_1_*/N*_2_) *≤τ*_1_, and *N*_2_ = ⟨*w*(*x*_1_)⟩ *τ*_1_ = *β* ⟨*x*_1_⟩ *τ*_1_ now is the signaling rate of the coupled reaction. Note that 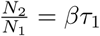 here. Compared to the decoupled bound (9), equation (12) contains one more term *Q ≥* 0 due to coupling. Note that this bound depends on the absolute value of *N*_2_ in addition to the relative signaling rate *N*_2_*/N*_1_.

This means the effect of noise suppression depends on both *N*_1_ = ⟨*x*_1_⟩ and *τ*_1_. To unravel this dependence, we first notice that if *τ*_1_ is small, then *Q ≈ τ*_1_ is small as well, so the bound (12) approaches the decoupled bound (9). Furthermore, since 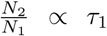, this means signaling rate is small in general, yielding 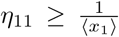, the Poisson noise lower bound. In other words, a small *τ*_1_ indicates that noise suppression below Poisson is infeasible. This is also intuitively clear, as small *τ*_1_ means *x*_1_ or plant dynamics is so fast that our controller is simply too slow to be effective.

If *τ*_1_ is not too small, then *Q* is of order *N*_2_. So whether coupling effect is significant depends on how *N*_2_ compares with 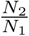. If *N*_1_ = ⟨*x*_1_⟩ is small, then relative signaling rate 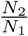 dominates while coupling has little influence, and the bound (12) approaches the decoupled bound (9) again, with minimum CV (i.e., 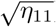) bounded by quartic root of 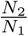. If *N*_1_ = ⟨*x*_1_⟩ is not too small, then *N*_2_ is comparable with 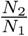 and coupling is advantageous. Indeed, in the other extreme where 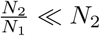, we have 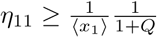, so the minimum CV is inversely proportional to the square root of *N*_2_ in contrast to *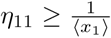* implied by the original Lestas *et. al.* bound, saying that minimum CV cannot be sub-Poisson. The above observation is consistent with example (6) in Section III-B, where small *n* corresponds to a *N*_2_ comparable with 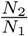, and large *n* corresponds to a *N*_2_ dominated by 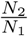.

In summary, from our lower bound (12) on noise suppression for coupled reactions, we see that coupling is most effective when *τ*_1_ is not too small and when absolute signaling rate *N*_2_ is at least comparable with relative signaling rate 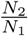. This suggests that coupling is most useful for energy efficient noise suppression, where 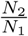 is not too large, so utilizing *N*2 through coupling to reduce noise further could be rather significant.

### D. Coupling in two-component CRN

Here, in order to discover design principles for noise suppression with coupling, we study the achievable lower bound for noise in two-node chemical reaction networks using LNA, explicitly allowing coupling among reactions.

We start from the following formula for noise for stable two-node chemical reaction networks using LNA (for derivation see Section V-B):

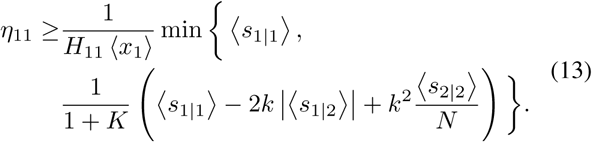

Here 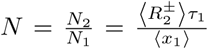, with *N*_1_,*N*_2_ being the signaling rates as in the Lestas *et. al.* bound in equation (9), 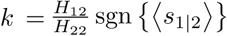, and 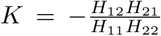. *k* is the controller actuation gain, as *H*_12_ is the logarithmic gain of *x*_2_’s influence on *x*_1_’s birth and death rates. The sign of ⟨*s*_1*|*2_⟩ enters *k* so that the sign of *k* is meaningful as well: *k <* 0 always implies amplification of noise, while *k >* 0 may result in suppression of noise. *K* is the closed loop gain, with negative sign in front to again make positive *K* correspond to noise suppression and negative *K*, noise amplification.

The two terms in the minimization in equation (13) are achievable (see Section V-A). *η*_11_ achieves the first term if the controller time scale is much larger than the plant time scale with *τ*_1_ ≫; *τ*_2_, while the second term is achieved when the controller time scale is much faster than the plant time scale with *τ*_1_ ≪ *τ*_2_. It is reasonable that *η*_11_ becomes 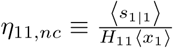 when the controller *x*_2_’s dynamics is very slow, as *x*_2_ is essentially constant in *x*_1_’s time scale, so *x*_1_ is doing birth-death by itself with constant *x*_2_. Therefore, we can regard *η*_11,*nc*_, the first term, as a baseline that is easily achieved when there is no control at all. Note that the no control case is not the same as open loop, as open loop could have controller actuation on the plant with no sensing.

The more interesting term is the second one, which could become less than the first no-control term in certain architectures. For the second term to be small, we need *k >* 0 and *K >* 0. *K >* 0 corresponds to negative feedback, with either *x*_1_ activating *x*_2_ and *x*_2_ repressing *x*_1_, or *x*_1_ repressing *x*_2_ and *x*_2_ activating *x*_1_. *k >* 0 corresponds to an actuation that is compatible with the coupling reaction. If we have positive coupling, i.e. ⟨*s*_1*|*2_⟩ > 0, then *k >* 0 implies *H*_12_ > 0 or that the controller *x*_2_ represses *x*_1_, as in example (4) of Section III-A. If we have negative coupling, i.e. ⟨*s*_1*|*2_⟩ *<* 0, then *k >* 0 implies *H*_12_ *<* 0 or that the controller activates *x*_1_, as in example (6) in Section III-B. More detailed implications are discussed below.

#### 1) Feedback vs. feedforward

Both incoherent feedforward and negative feedback architectures are widely found in natural biological regulatory systems with homeostasis [29]. However, our result shows that feedforward architectures always have larger noise compared to negative feedback architectures with the same plant dynamics, controller actuation, and coupling. Feedforward architectures allow controller actuation but no sensing, so *K* = 0. If *K >* 0, which is the case for negative feedback, then the lower bound for noise is always lower than the feedforward case by a factor of 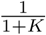. Note that noise could be amplified by positive feedback with *K <* 0.

This is in strong contrast with the result for deterministic robust perfect adaptation, relevant for suppression of extrinsic noise (i.e. noise of parameters). Robust perfect adaptation is the property that a deterministic system could reach a homeostasis at steady state despite constant disturbances and uncertain dynamics [30], which is biologists’ term for robust constant disturbance rejection. Both feedforward and feedback architectures could implement robust perfect adaptation equally well, with only differences in the physical controller implementation considerations [31]. Once noise is considered, we see that negative feedback could achieve smaller noise than feedforward, even when the underlying deterministic system dynamics is the same.

#### 2) Coupled vs. decoupled

The coupled case always have lower noise than the decoupled case if *k >* 0. If we assume our system does not have coupling, then ⟨*s*_1|2_⟩ = 0, so the term –2*k* |⟨*s*_1*|*2_⟩|, which is negative when *k >* 0, becomes zero for the decoupled case.

We also see that larger *N* or higher signaling rate always corresponds to less noise in the decoupled case, which agrees with what is implied by the Lestas *et. al.* bound. The lower bound for this case is 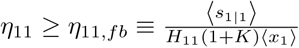, achieved in the large signaling (*N* → ∞) and fast controller (*τ*_2_ ≪ *τ*_1_) limit. Therefore we can regard *η*_11,*fb*_ as the lowest noise achievable with only feedback regulations and no coupling. Note that 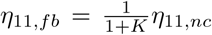, so we decrease noise by a factor of 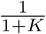 compared to no control case when we have negative feedback.

In comparison, the coupled case have a different interpretation of the relative signaling rate *N*, as 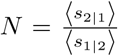 if ⟨*s*_1|2_⟩ ≠ 0, a consequence of equation (3). In this case, the relative signaling rate and the coupling strength are strongly related to each other. In particular, if the coupling reaction is known, then the optimal signaling rate is no longer infinite. This is shown explicitly in the following subsection.

#### 3) Limit on noise suppression by coupling

We show that the relative signaling rate *N* is not larger the better for coupled systems and that the suppression of noise through one coupled reaction is limited to a factor of 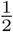 beyond the feedback lower bound *η*_11,*fb*_.

Consider the case where we have only one coupling reaction of the form {*x*_1_, *x*_2_} → {*x*_1_+ *b*_1_*σ*_1_, *x*_2_ + *b*_2_*σ*_2_}, where *b*_1_, *b*_2_ *∈* ℤ_>0_ are the molecule counts of *x*_1_ and *x*_2_ changed by this reaction, and *σ*_1_, *σ*_2_ *∈ {*1, *-*1*}* are signs of the change. We assume this coupled reaction has *α*_1_ portion of the birth or death flux of *x*_1_, and *α*_2_ portion of the flux of *x*_2_, where *α*_1_, *α*_2_ *∈* (0, 1]. We also assume that all reactions other than the coupled one have step size 1. In terms of these parameters, we have 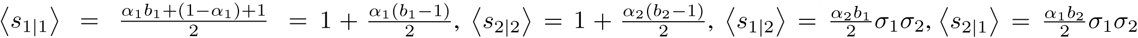, and 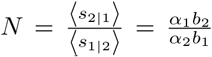. Substituting these into equation (13) yields lower bounds for noise in terms of *α*_*i*_ and *b*_*i*_. Notice that once the coupling reaction is known or *b*_1_, *b*_2_ are determined, varying the relative signaling rate *N* is the same as varying *α*_1_ and *α*_2_, the fractions of fluxes that the coupled reaction take.

The partial derivative of the second term of the lower bound in equation (13) with respect to *b*_1_ is always positive, so *b*_1_ should be as small as possible, while the partial derivative with respect to *b*_2_ is always negative, so *b*_2_ should be as large as possible. This makes sense, as larger 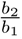 means that *x*_1_’s change is amplified in *x*_2_’s change, so the signal is less corruptible by internal noise of *x*_2_.

If we take *b*_1_ = 1, then the partial derivative with respect to *α*_1_ is always negative, so *α*_1_ = 1 is optimal. This is reasonable, as *α*_1_ = 1 means the coupling reaction accounts for as large a fraction of change as possible of *x*_1_, so the knowledge of *x*_1_’s change obtained from *x*_2_ through the coupling reaction is more informative.

The partial derivative with respect to *α*_2_, however, is usually not as simple. The optimal *α*_2_ is 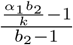 when *b*_2_ ≠ 1. This shows that the optimal fraction of flux in *x*_2_ of the coupled reaction is related to the controller actuation gain as well as the stoichiometry of the coupled reaction. In particular, this shows that the optimal relative signaling rate *N* is determined by parameters of the coupled reaction, such as *b*_1_, *b*_2_ and *k*. For example, taking *α*_1_ = 1 and large *b*_2_, we have 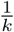 as the optimal *α*_2_, and the optimal effective signaling rate *N* is 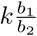.

It should be noted that the relative signaling rate is not larger the better when the coupled reaction is fixed does not contradict that larger relative signaling rate should always reduce noise as indicated in equation (13). Indeed, larger effective signaling rate *N* always corresponds to smaller lower bound for noise if we do not constrain *b*_1_ and *b*_2_. The optimal *b*_2_ is infinity, which corresponds to infinite *N*. However, once *b*_1_ and *b*_2_ are fixed, then *N* is no longer free to vary by itself. For example, if we fix parameters at their optimal values so that *b*_1_ = 1, *α*_1_ = 1 and *b*_2_ is large, then 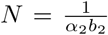, which could only become larger by decreasing *α*_2_. However, smaller *α*_2_ would also mean that the coupling reaction accounts for a smaller fraction of *x*_2_’s change, so the signal about *x*_1_ becomes more easily obfuscated by noise of *x*_2_’s other reactions not coupled with *x*_1_. In other words, since smaller *α*_2_ corresponds to more signal amplification through *x*_2_’s own birth-death reactions that are not coupled with *x*_1_, there is a tradeoff between noisy amplification of signals through the decoupled reactions and the preservation of accurate signals through the coupled reactions.

Another implication of the above observation is that with coupled reactions, a moderate signaling rate could result in significant noise suppression, as the optimal *N* could be small. This suggests that noise suppression with coupling could be rather cost-effective, therefore preferable for biological systems.

Now, taking the optimal values of *b*_1_ = 1, *α*_1_ = 1, *b*_2_ *→ ∞*, and 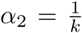 results in 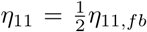 in the fast controller limit (*τ*_2_ ≪ *τ*_1_), so the optimal lower bound with one coupling reaction while other reactions have step size 1 is half of the optimal lower bound of negative feedback. This shows that coupling cannot make noise arbitrarily small beyond the decoupled case.

## IV. Discussion

In this work we investigated the coupling mechanisms that is ubiquitous in biology for noise suppression. We constructed examples that can suppress noise below the Lestas *et. al.* bound [2] and, by systematically analyzing all two-node chemical reaction networks, showed that coupling has implications on effective network architectures and that coupling has its own limitations in noise suppression.

Our example constructed in Section III-B suppresses noise below the Lestas *et. al.* bound, but broke one of their assumptions that require the plant and the controller to have separate birth-death reactions. It may therefore seem natural that we could go below their bound. However, it should be noted that not only is the Lestas *et. al.* bound hypothesized to be true in much more general situations [2], previous efforts in going beyond this bound either employed complex non-biological controllers [32] or could only show a theoretically better lower bound with unknown achievability [33]. Hence, finding an example or a class of systems that suppresses noise beyond the Lestas *et. al.* bound is highly nontrivial with significant implications. In particular, the example in Section III-B shows that biologically plausible reaction systems could indeed suppress noise below the Lestas *et. al.* bound, and situations beyond the bound should be seriously explored and considered for regulation and information processing in biological systems.

With that said, the results in this work does not try to contradict the results in [2], but rather follows its pioneering efforts in exploring fundamental limits of noise suppression in biological systems. We showed that noise suppression with coupling could further suppress noise, but still has its own fundamental limitations. This is also connected with the work of [3], which does not assume LNA and derives tight exact lower bounds for noise, but the lower bounds are harder to analyze for different cases. It is therefore of high interest to derive exact lower bounds in the spirit of [3] that includes coupling. Furthermore, theoretical investigations that relate LNA analysis with the methods of [3] would be highly beneficial, as the LNA method is scalable and analytically tractable, hence useful for controller design and architecture exploration.

This work focused on intrinsic noise, while noise in biology could be intrinsic as well as extrinsic. Intrinsic noise arises from stochastic nature of the system itself, while extrinsic noise comes from fluctuations in system parameters [34]. Extrinsic noise could be treated rather well by ignoring intrinsic noise [34], so perfect extrinsic noise suppression could be considered as perfect adaptation in the deterministic system. Robust perfect adaptation and its constraints on system architecture for biomolecular systems is studied in [31]. It is therefore desirable to connect intrinsic noise suppression together with robust perfect adaptation concerns. After all, noise suppression is not meaningful if the fixed point is not desirable.

While coupling considered in this work is of a binary kind, where two components are either coupled or decoupled, several biological phenomena observed in nature also have a “soft” coupling interpretation. For example, genes that are spatially close to each other due to their position on the genome or its 3D structure are more likely to be transcribed together in a short period of time [8]. Indeed, the positioning of genes are evolutionarily selected and the 3D structure of the genome is highly regulated [12]. This urges the study of a generalization of the coupling notion here to include these cases. It is also worth investigating how these general “soft” coupling notions relate to the cases investigated in Section III-D where coupling is only a fraction of the birth-death fluxes.

Lastly, coupling is a phenomenon rather specific to biomolecular control. Although covariance is considered in canonical stochastic control theory, coupling is a structually and physically different way of influencing variables’ covariance than regulations through birth and death rates. This echoes the results in [31], where physical constraints specific to biomolecular reaction systems is shown to make integral feedback implementation a nontrivial design problem in biology. With these observations, it is highly desirable to develop theoretical tools custom-fit for biological systems so as to gain insight for biomolecular control problems.

## ACKNOWLEDGEMENT

The authors would like to thank Noah Olsman for constructive discussions. F. X., J. Y. and J. C. D. are partially funded by the Defense Advanced Research Projects Agency (Agreement HR0011-16-2-0049). J. Y. acknowledges the support of NSF/Simons Foundation. The content of the information does not necessarily reflect the position or the policy of the Government, and no official endorsement should be inferred.

### V. Appendix

#### A. Two component CRN noise analysis

Here we derive equation (13) using LNA.

Equation (2) for the two dimensional (*n* = 2) case gives rise to the following system of equations:

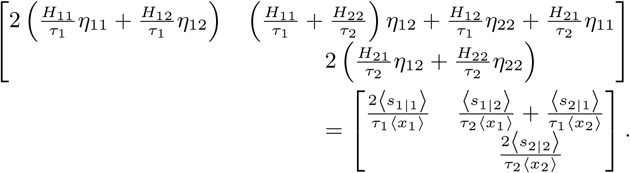

We make the following assumptions: the deterministic linearized system is stable, and *H*_11_, *H*_22_ ≠ 0. The first assumption is needed for the LNA to have finite variance. Since the deterministic system is stable if and only if –***M*** is Hurwitz, for *n* = 2 case we have the following conditions for stable linearized system: 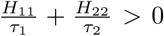 while *H*_11_*H*_22_*- H*_12_*H*_21_ > 0. The second assumption is more biological. *H*_11_ = 0 implies 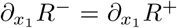, so stability of *x*_1_ relies on sensing and actuation of other system components. This is highly undesirable biologically, as constantly mutations could occur in the system that results in loss of function, breaking the sensing reactions such that other components become fixed and do not respond to changes of *x*_1_, or breaking the actuation reactions such that other components’ actions on *x*_1_ become zero. In either case, *x*_1_ dynamics could become unstable, implying disastrous consequences for the biological organism. In addition, the *H*_11_ = 0 or *H*_22_ = 0 cases are not biologically informative but simple to analyze, and can be easily shown that they satisfy the bound (13) as well.

With the above assumptions, we could then write the equation above into more informative parameters and rewrite it into the following linear system of equations:

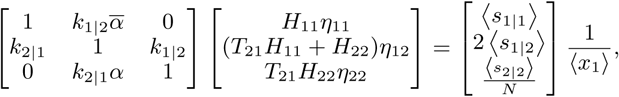

where 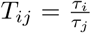 is a ratio of time scales, 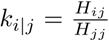 is the logarithmic sensitivity of *x*_*i*_ to changes in *x*_*j*_ normalized by the sensitivity of *x*_*j*_ to its own changes, 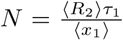 is the relative signaling rate as defined in [2], and 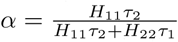 with 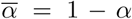. Solving the system for *η*_11_ yields the following:

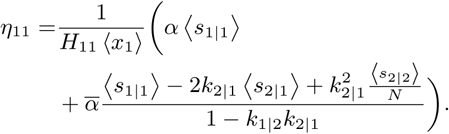

So we see that *η*_11_ can be expressed as a convex combination of two terms since 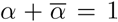 and *α ∈* [0, 1]. Taking the min of the two terms then yield the lower bound shown in equation (13), with the first term achieved when *α* = 1 or *τ*_2_ ≫ *τ*_1_, and the second term achieved when *α* = 0 or *τ*_2_ ≪ *τ*_1_.

#### B. Birth-death interpretation of LNA

LNA was considered an interpretable but inaccurate approximation because of its derivation assuming large mean and Gaussian noise, which violates the non-negativity and discreteness of the variables [26], [22]. Here, through one simple example, we show that the Gaussian interpretation is not necessary, and the non-negativity and discreteness could be preserved. A systematic investigation is ongoing work to be published on another occasion.

Consider the following non-linear birth death process:

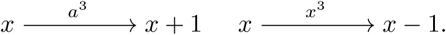

Equation (2) gives 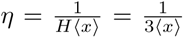. The only fixed point is *x*^*∗*^ = *a*. Then consider the following linear-propensity birth-death process:

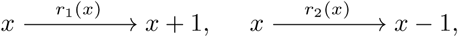

where 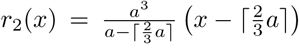 and *r*_1_ (*x*) = *r*_2_ (*x*) *-* 3*a*^2^(*x - a*), for *a >* 3. Therefore *R*^*-*^ *- R*^+^ for this system is the same as that for the original nonlinear birth-death process, and equation (2) would yield the same result 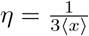 with the same fixed point *x*^*∗*^. On the other hand, the system does not evolve below 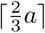 as the death rate is zero on that number, so the variable *x* is always non-negative.

This implies that LNA could be a good approximation even when the mean molecular count is small (in fact as small as 3 in this case), and the distribution could preserve the discreteness as well as non-negativity, so it is not necessarily Gaussian. The only situations that LNA breaks down then are the same as the situations where the deterministic linearization of a nonlinear dynamical system breaks down: sharp non-lineariarities away from the fixed point.

#### C. Channel capacity of coupled reactions

The key step to derive the lower bound of noise suppression with or without coupling is the find the lower and upper bounds of the mutual information between *x*_1_ and *x*_2_. Here we first introduce the mathematical definition of mutual information between two random processes (rather than variables), and compute the channel capacity of a coupled reaction.

Consider a complete probability space (Ω, *ℱ*, P) with nondecreasing right continuous family of sub-*σ* algebras (ℱ_*t*_), *t ϵ* [0, *T*]. Let (*n*_*t*_, *ℱ*_*t*_) be a Poisson process with its intensity *λ*_*t*_ being a function of of the input signal *θ* = {*θ*_*s*_, *s ⩽t*} and also *n* = {*n*_*s*_, *s ⩽* _*t*_} in an non-anticipatory manner:

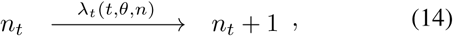

where the non-anticipatory constraint requires that *λ*_*t*_(*t, θ, n*) is *ℱ*_*t*_-measurable. If a certain coding *λ*_t_ (*t, θ, n*), 0 *⩽t ⩽T* is given, then a natural question is how much information is contained in the received signal *n* about the transmitted signal *θ*. By definition, the mutual information is:

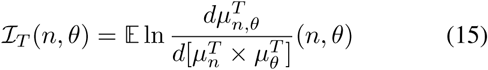

where 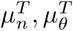 and 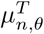 are measures corresponding to processes *n*_*t*_, *θ*_*t*_, and (*n*_*t*_, *θ*_*t*_), 0 ⩽*t ⩽T*.

The Liptser-Shiryaev formula shows that [35]:

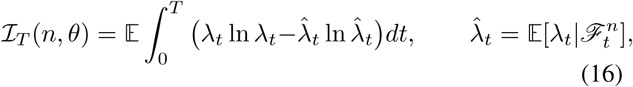

immediately we can see if *λ*_*t*_ is *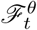*-measurable, then ℐ_*T*_ (*n, θ*) = 0 since *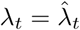*

By the causality nature of such channel, the Liptser-Shiryaev formula is actually the formula for directed information: ℐ_*T*_ (*θ → n*) (see [36] for a complete introduction of directed information). Here we show that if we have one another parallel channel:

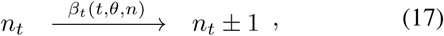

where *β*_*t*_(*t, θ, n*) is again ℱ_*t*_-measurable, then the mutual information (or more precisely, the directed information) from *θ*_*t*_ to *n*_*t*_ is the sum of mutual (directed) information of each channel:

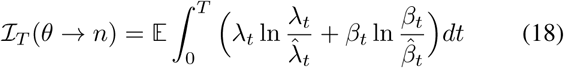

Equation (18) can be bounded by using Lagrangian multiplier methods, assuming the first and second moments of the propensities are finite, therefore we obtain an upper bound of the mutual information of two parallel Poisson channels (equation (14) and (17)) [2]:

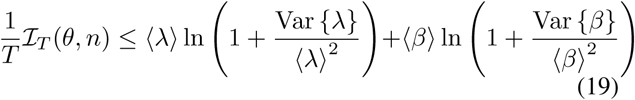

Now let’s turn to the coupled reaction, using equation (11) as an example. Equation (11) is equivalent to two Poisson channels:

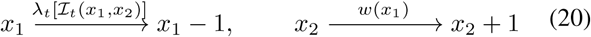

where *λ* switches between 0 and a large constants *K*. It switches from 0 to *K* when *x*_2_ increases by 1, then switches back to 0 when *x*_1_ also increases by 1. Hence the overall effect of the first reaction to tracking the dynamics of the second reaction, when the second reaction occurs, the first immediately occurs, when setting *K → ∞*. Then the channel capacity of this coupled reaction can be calculated by summing up the channel capacity of the two channels (equation (20)). The second one, as the same as a usual Poisson channel, has a channel capacity *C*_2_ = Var {*w*(*x*_1_}*/ ⟨w*(*x*_1_) ⟩. For the first reaction, since *λ* basically tracks the number of occurrence of the second one, we instead can use a classic conclusion from large deviation theory, which claims that Var {*λ*} is equal to its mean [37]. On the other hand, in the limit of long time, the empirical mean of number of events is equal to ⟨*w*(*x*_1_) ⟩, then from ergodic theorem we conclude that ⟨*λ⟩* = ⟨*w*(*x*_1_) ⟨. Therefore the channel capacity of the coupled reaction is then:

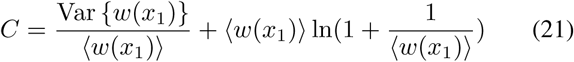

Furthermore, coupling does not alter the lower bound on mutual information, which is derived from Pinsker’s non-anticipatory *ε*-entropy [33], [38]. So the lower bound on noise suppression is only due to an increase of channel capacity.

